# DRP1-mediated mitochondrial dynamics orchestrate EMT in glioblastoma cells

**DOI:** 10.64898/2025.12.12.694080

**Authors:** Mahima Choudhary, Megha Chaudhary, Nargis Malik, Kunzang Chosdol, Swarnima Kushwaha, Anirudha K Sahu, Shibasish Chowdhury, Bhawana Bissa, Rajdeep Chowdhury, Sudeshna Mukherjee

**Affiliations:** Dept. of Biological Sciences, Birla Institute of Technology and Science, Pilani, Rajasthan, India; Dept. of Biochemistry, Central University of Rajasthan, Ajmer, Rajasthan, India; Dept. of Biochemistry, All India Institute of Medical Sciences, Delhi, India

## Abstract

**Background:** Epithelial to mesenchymal transition (EMT), a differentiation process, frequently imparts invasive properties in Glioblastoma Multiforme (GBM), which leads to a poor prognosis. Cells lose apical-basal polarity, cell-cell connections, and/or chemo-resistance during EMT, which can result in the spread of cancer and the acquisition of additional stem cell-like traits. It is unclear how organelle dynamics influence EMT in this respect. The interaction between cytoskeletal and mitochondrial regulators governing GBM cell EMT is explored in this article.

**Results and Discussion:** In GBM cells, we observed that TGF-β-induced EMT led to a proliferative arrest, which was accompanied by a fragmented mitochondrial morphology, elevated expression of fission markers such as DRP1, MFF, and FIS1, and most importantly, localization of mitochondria near the cell boundaries. An increase in mitochondrial ROS accompanied this, but their functional status was indicated by a higher oxygen consumption rate (OCR). Additionally, cytoskeleton re-distribution and EMT reversal were the outcomes of si-RNA-mediated elimination of the fission-marker DRP-1 or pharmacological inhibition of fission by Mdivi-1. On the other hand, drugs that disrupt the cytoskeleton, shifted the spatial distribution of mitochondria to the perinuclear area, which had an adverse effect on EMT. Notably, it was shown that RhoA, a protein that helps organize the actin cytoskeleton, co-immunoprecipitates with DRP1 and governs both cytoskeletal dynamics and mitochondrial fission in its presence.

**Conclusion:** Our research sheds substantial insight on the current interactions between the cytoskeleton and mitochondrial spatial dynamics that control EMT in GBM cells, which may have significant therapeutic implications.

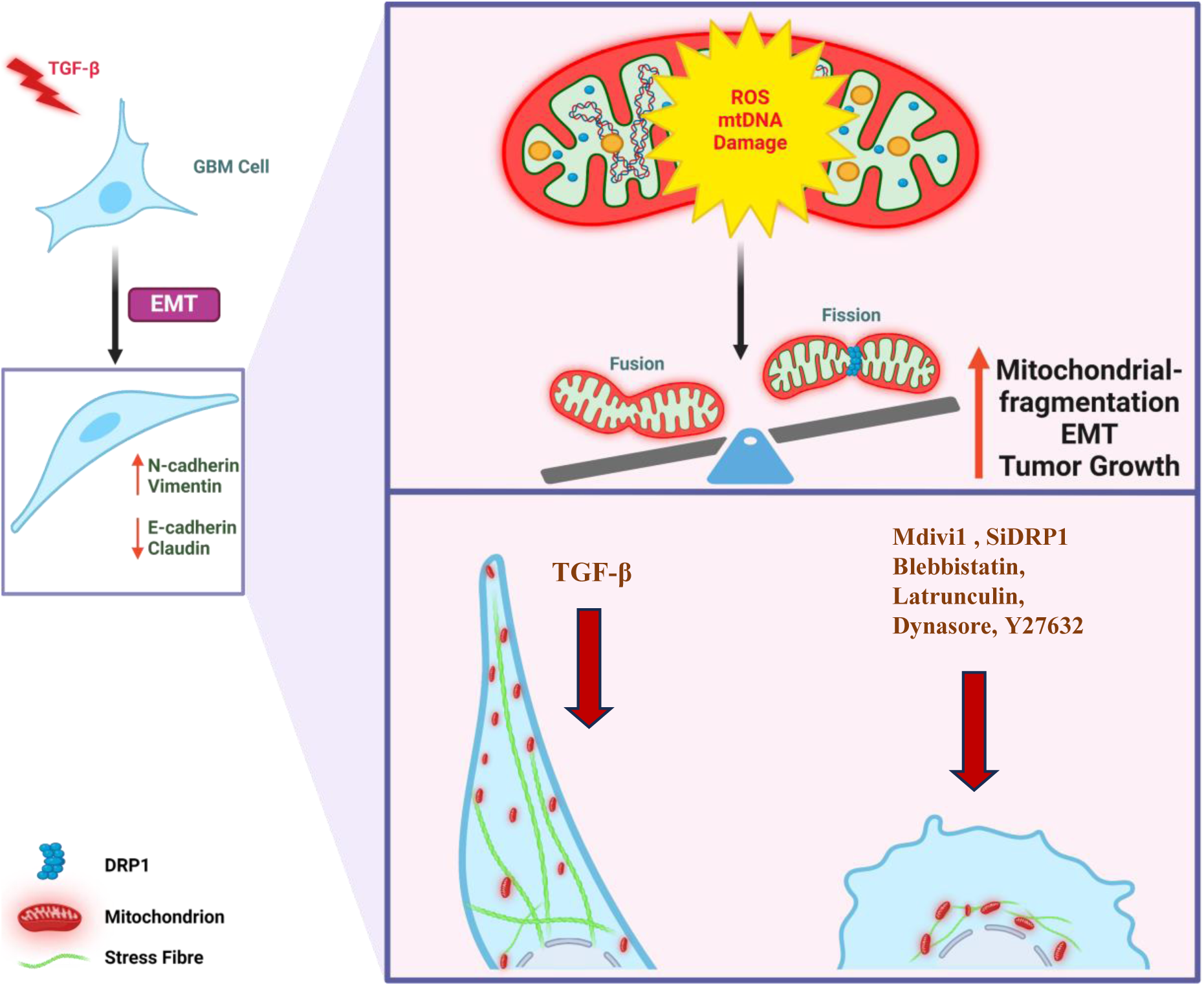

## Introduction

Glioblastoma multiforme (GBM) represents the most common and aggressive primary malignant tumor of the central nervous system (CNS) in adults. It accounts for roughly 57% of all gliomas and 48% of primary malignant tumors, with a median survival of 12 to 18 months [1], [2]. Characterized by its highly infiltrative nature, rapid progression, and resistance to treatment, GBM poses a significant challenge to the effective clinical management of the disease [3]. Importantly, even after successful surgery, GBM often recur intracranially and/or develop intracranial metastases. In this context, extracranial metastasis, though rare, patients have an average overall survival period of only 17 months, underlining the importance of therapeutically addressing the aggressive and invasive characteristics of GBM cells. Collectively, both intra and extracranial metastases pose a significant risk to the long-term survival of GBM patients[4], [5].

Herein, the intricate interplay between the tumor cells and the tumor microenvironment (TME) further exacerbates the aggressive behavior of GBM. The TME comprises a complex milieu of immune cells, stromal cells, extracellular matrix components, and, most importantly, soluble factors, which can potentially influence tumor growth, invasion, and therapeutic response [6]In this regard, one of the significant cytokines predominantly prevalent in GBM TME and driving GBM progression is the transforming growth factor-beta (TGF-β) [7]. This cytokine’s classical function in regulating tumor pathophysiology involves triggering the tumor cells’ epithelial-to-mesenchymal transition (EMT) through SMAD-dependent or SMAD-independent pathways. Herein, EMT is a fundamental process adopted by embryonic cells for migration; however, when dysregulated, it can contribute significantly to the invasive behavior of the tumor cells. Therefore, understanding the regulation of EMT is critical to harnessing the invasive characteristics of GBM cells [8].

In this context, TGF-β is known to dynamically regulate the cytoskeleton, enabling an increased migratory and invasive capability to facilitate the metastatic escape of the tumor cells. This might include re-arrangement of matrix adhesions, differential actin-myosin contractility, and further proteolytic remodeling of tumor tissues to promote EMT and subsequent steps of the metastatic cascade [6], [9]. However, inculcation of this re-programmed behavior in tumor cells is energy expensive, and therefore, tumor cells must dynamically modify their energy production and consumption to local conditions meeting their invasive demands. Herein, tumor cells have this astounding ability to regulate mitochondrial dynamics to meet their local energy and metabolic needs [10], [11], [12]. These dynamic ATP-producing organelles can alter in size and location within cells through fission and fusion cycles. The highly conserved dynamin-related proteins (DRPs) control the opposing processes of mitochondrial fission and fusion. For example, mitofusins (Mfn) 1 and 2, also members of the DRP family, perform outer membrane fusion, while the mitochondrial fission protein Drp1, facilitates mitochondrial outer membrane scission [11], [13]. Simultaneously, mitochondria also rely on the cytoskeleton for their intracellular movement, which is essential for meeting the local cellular energy needs [10], [14], [15]. Thus, the coordinated interaction between mitochondria, motor proteins, and cytoskeletal tracks facilitates efficient transport, distribution, and anchoring of mitochondria within the cell, thereby contributing to adaptation to changing metabolic demands and altered cellular physiology[16], [17], [18], [19]. In this context, the link between mitochondria, cytoskeleton, and EMT in GBM cells is poorly explored. Therefore, understanding the molecular mechanisms underlying their reciprocal regulation may offer novel insights into the pathophysiology of EMT-associated diseases like GBM. It may also significantly transform the development of targeted therapeutic strategies for invasive cancers.

## Results

### TGF-β results in Smad-dependent EMT in GBM cells

GBM is a highly invasive tumor with correlated morbidity and mortality. To approximate the invasive features of GBM cells, we analyzed molecular markers associated with a key cellular event promoting invasion and metastasis of the tumor cells, which was epithelial to mesenchymal transition (EMT). Glioblastoma patient samples were analyzed for expression of EMT markers. They showed a significant up-regulation of proteins-Vimentin (>20 fold) and N-cad (>4 fold) as compared to normal brain, highlighting their intricate involvement in EMT and glioma progression **(Figure 1A)**. Furthermore, the human protein atlas data analysis also revealed that the EMT markers like Vimentin and N-cad are important for GBM prognosis. Their elevated levels are associated with poor survival of GBM patients (**Figure 1B).** Gene Ontology (GO) analysis of differentially expressed genes from GBM patient data and mapping of their corresponding pathways further revealed de-regulation of cytoskeletal and cellular differentiation-associated processes **(Supp. Fig 1A)**. These observations pinpointed the importance of further analyzing the molecular regulation of cytoskeleton-driven cellular differentiation processes like EMT in GBM pathogenesis. Therefore, to induce an EMT-like invasive phenotype, we exposed the GBM cells to the classical EMT-inducing cytokine-TGF-β. It is known to be predominantly present in the GBM microenvironment and plays a profound role in GBM progression. As expected, TGF-β significantly enhanced the expression of Vimentin and N-cad in all three GBM cell types studied-U87MG **(Figure 1C, D)** and U373, and LN229 **(Supp. Figure 1B**). The change in molecular expression of EMT markers was also associated with a change in cellular morphology as evident from the SEM images of U87MG reflecting an extended filamentous cellular phenotype, a hallmark of EMT (**Figure 1E**). Herein, earlier studies suggest that cells undergoing EMT can re-program major cellular functions including metabolism and proliferation to attain the differentiated phenotype [20], [21]. Therefore, we characterized the allied effects of EMT on the GBM cells to better understand the specific key regulators governing this complex transdifferentiating process. Notably, while our GBM cells acquired a phenotypic transition, TGF-β exposure had a minimal effect on GBM cell death, as observed through dose and time kinetic study in GBM cell lines (**Supp. Figure 1C**); however, interestingly a transition from epithelial to mesenchymal state was associated with a cellular growth arrest in GBM cell lines (**Supp. Figure 1D).** In addition, an increase in the expression of cyclin-dependent kinase inhibitor-p21 and a simultaneous decrease in levels of the proliferation marker-proliferating cell nuclear antigen (PCNA) was observed corroborating an attenuated proliferative state in U87MG cell line (**Supp. Figure 1E**). TGF-β is known to function through canonical SMAD-dependent and/or non-canonical SMAD-independent pathways. Herein, an increased expression of SMAD2 and p-SMAD2 was observed upon TGF-β treatment in U87MG cell line (**Figure 1F)**. Inhibition of TGF-β signaling using SIS3 (pharmacological inhibitor of SMAD) or SB431542 (TGF-β receptor inhibitor) led to a reversal of associated EMT-like features in U87MG cell line like round or more flattened cell morphology **(Supp. Figure 1F and Figure 1G**). Finally, this complex re-programming involving change in cellular shape and dimension, inhibition of DNA replication, and growth arrest was also associated with an enhanced intracellular ROS in U87MG cell line (**Figure 1H**). The above studies indicate that the GBM cells undergoing EMT regulate several cellular circuitries to achieve this transition.

**Figure 1.**
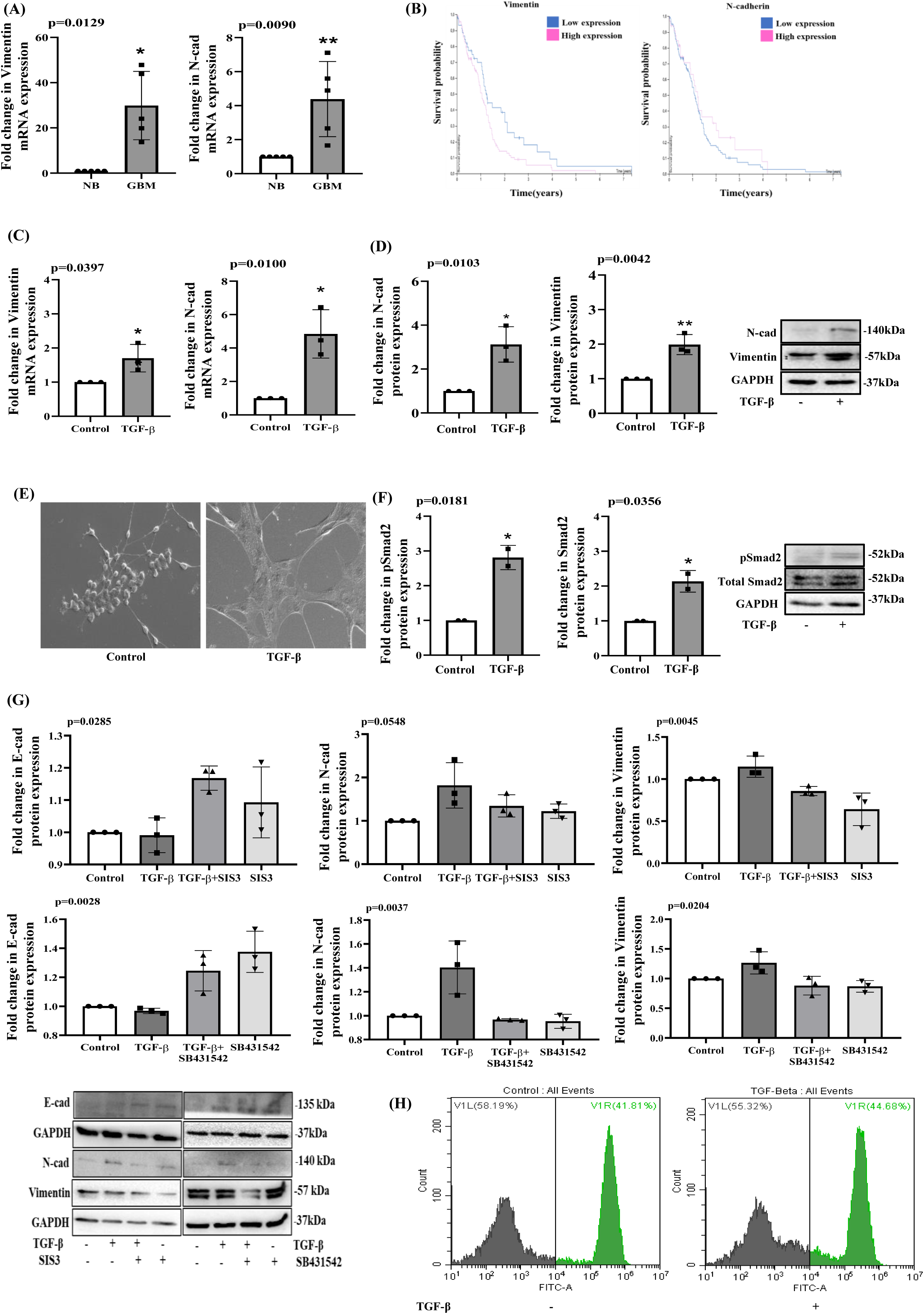
TGF-β results in Smad-dependent EMT in GBM cells: **(A) Bar graph represents the** change in mRNA expression of Vimentin (p=0.0129) and N-cad (p=0.0090) of normal vs GBM patient RNA samples. (**B)** Kaplan-Meier plot obtained from human protein atlas correlates Vimentin and N-cad expression with survival of GBM patients. The blue and pink curves represent low and high protein expression, respectively. **(C)** The bar graph represents changes in Vimentin (p=0.0397) and N-cad (p=0.0100) mRNA expression in U87MG cells treated with TGF-β with respect to control cells. (**D)** Bar graph and immunoblot represent changes in N-cad (p=0.0103) and Vimentin (p=0.0042) protein expression in U87MG cells treated with TGF-β with respect to control cells. GAPDH serves as a loading control. **(E)** Scanning electron microscope (SEM) images of U87MG cells after treatment with TGF-β (20ng/ml) for 72 hours; Scale bar 50μm. **(F)** The bar graph and immunoblot represent changes in expression levels of phospho-Smad2 (p=0.0181) and Smad2 (p=0.0356) in U87MG cells. **(G)** Bar graphs and immunoblots represents changes in E-cad(SIS3;p=0.0045 & SB431542; p=0.0204), N-cad(SIS3;p=0.0548 & SB431542; p=0.0037) and Vimentin(SIS3;p=0.0285 & SB431542; p=0.0028), protein expression upon inhibiting Smad using SIS3 (10µM) and TGF-β receptor inhibitor SB431542 (10µM) alone or in combination with TGF-β for 72 hours in U87MG cells. **(H)** Cellular ROS was measured using DCFDA through flow cytometric analysis in U87MG cells. The results of our study are represented as mean ±SD *(n*=3). Paired, two-tailed t-tests were used, where (*) p<0.05, to determine the significance when two groups were compared.

### Structural and distributional changes of mitochondria are associated with TGF-β exposure

Cellular oxidative metabolites like ROS play a critical role in the occurrence and development of gliomas alongside modulation of the GBM microenvironment. A high or lower than threshold level of ROS can potentially regulate growth-related signaling pathways [22]. In this regard, recent studies also couple ROS-induced growth arrest with mitochondrial de-regulation [23]. Thus, it is evident that the cell cycle and mitochondrial homeostasis are coordinated and interdependent; however, the interdependence between ROS, growth arrest- and mitochondrial functions during EMT remains largely unexplored. Herein, we hypothesized that, since mitochondria are the hub for ROS generation and mitochondrial ROS (mtROS) signaling can potentially facilitate cellular homeostasis, they might be critical to EMT re-programming in GBM cells. In correlation to the above, we observed that TGF-β exposure resulted in a significantly increased production of mtROS in both U87 and U373 (**Figure 2A & Supp. Figure 2A**) coupled to a mitochondrial membrane depolarization in U87MG cell line (**Figure 2B**). In accordance with, an increase in total ROS and associated cell viability was also observed in U87MG cell line (**Supp. Figure 2B).** Interestingly, this alteration in mitochondrial state was linked to an increased localization of mitochondria towards the cell periphery, enhanced mitochondrial circularity, increased mitochondrial fragmentation, and simultaneous reduction in mitochondrial perimeter suggesting an exhaustive overhaul of mitochondrial homeostasis during EMT transformation in U87MG and U373 cell line respectively (**Figure 2C & Supp. Figure 2C**). The altered mitochondrial distribution was validated through immunofluorescence staining of outer mitochondrial membrane protein TOM20 which showed similar results in U87MG cell line (**Figure 2D**). Herein, it is well-established that mitochondrial dynamics is maintained through fission and fusion cycle regulating mitochondrial quality control and energy production. In this context, dynamin-related protein 1 (Drp1) is a GTP-hydrolyzing enzyme that catalyzes cellular mitochondrial fission by binding to the outer mitochondrial membrane (OMM); the Drp1 receptor-MFF enables the binding of the hydrolyzing enzyme, and FIS1 involves in fission functions by regulating assembly of Drp1-complex[24]. Interestingly, our study observed an increased expression of DRP1, FIS1, and MFF coupled to the EMT-associated altered distribution of mitochondria in the U87MG cell line **(Figure 2E and F)**. This was further validated as we observed an βincreased level of DRP1 and phospho-DRP1 (pDRP-1; Ser616), which is known to promote the localization of DRP1 to the OMM in U87MG cell line **(Figure 2F and G)**. To further understand the interdependency of ROS and mitochondrial fission, we quenched ROS with N-acetylcysteine (NAC). This resulted in a decrease in the level of DRP1, suggesting that ROS generation facilitates the fission event in U87 cell line **(Figure 2H)**. Mitochondrial fission is often linked to the compromise of mitochondrial function and subsequent cellular apoptosis [25]. However, as mentioned earlier, we failed to observe a significant cell death whilst cells underwent EMT during the time period of our study. However, as we analyzed the mitochondrial oxidative phosphorylation (OXPHOS), which is a key energy-producing pathway often utilized by tumor cells to maintain cellular harmony [26], [27]; interestingly, we observed a high Oxygen Consumption Rate (OCR) in GBM cells upon ββ treatment indicative of re-distributed, yet functional mitochondria in U87MG cell line **(Figure 2I).**

**Figure 2.**
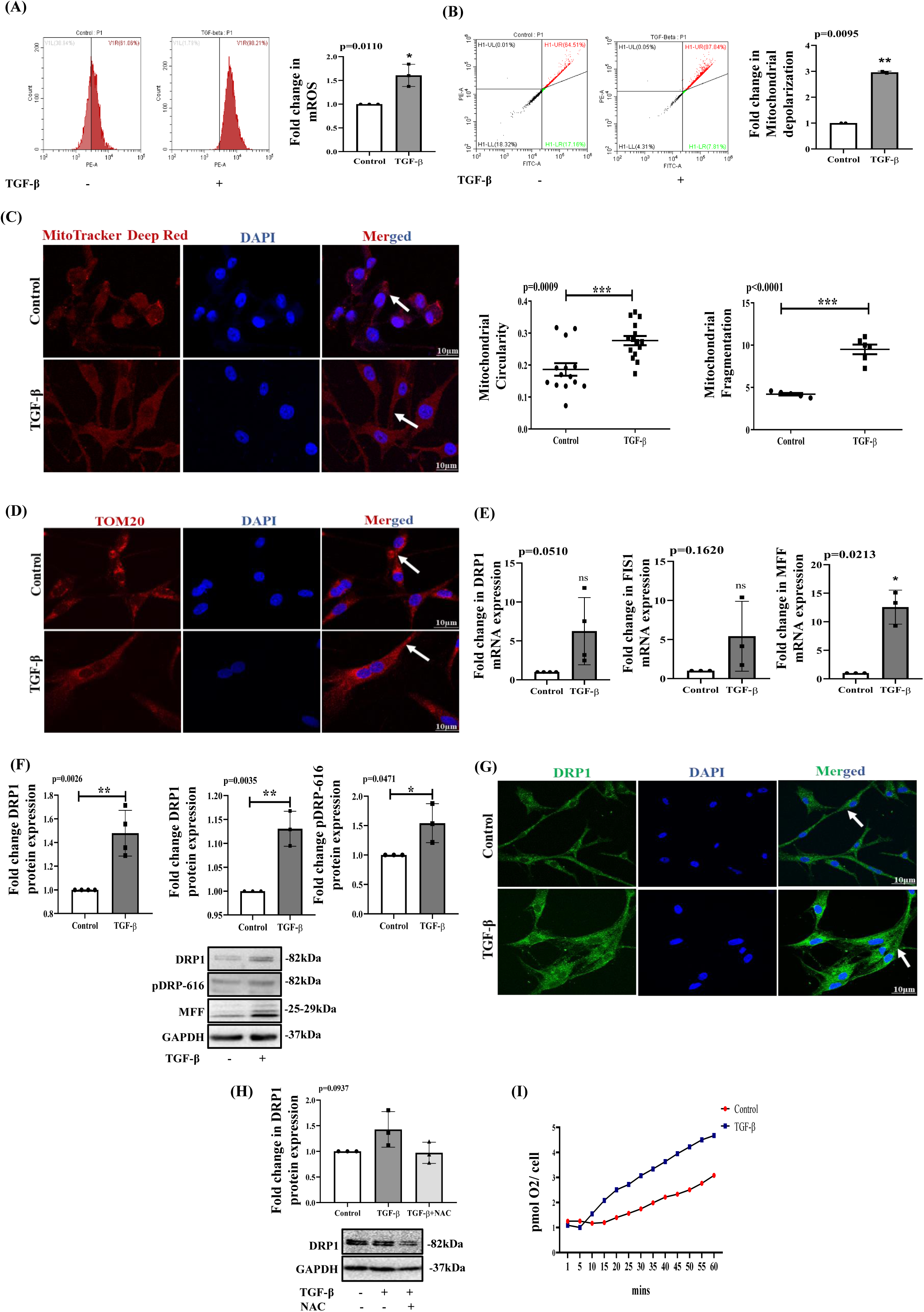
Structural and distributional changes of mitochondria are associated with TGF-β exposure: **(A)** Mitochondrial ROS measured using MitoSOX Red post-TGF-β treatment through flow cytometric analysis represented a shift in mtROS generation in U87MG cell line (p=0.110). **(B)** Analysis of mitochondrial potential by JC-1 post TGF-β treatment for 72 hours (p=0.0095) in U87MG cells. **(C)** Mitochondrial staining was performed on U87MG cells using MitoTracker Deep Red, and DAPI was used to stain the nucleus (shown in blue). Quantification was done using ImageJ software. Scale bar: 10µm and the bar grapgh represents the fold change in mitochondrial circularity (p=0.0009) and fragmentation (p<0.0001). **(D)** Immunofluorescence images representing TOM20 expression and localization post-TGF-β treatment for 72 hours in U87MG cells. The nucleus stained with DAPI is shown in blue. Scale bar: 10µm. **(E)** Changes in mRNA expression of mitochondrial fission-related genes such as DRP1(p=0.0510), FIS1(p=0.1620), and MFF(0.0213) post-treatment with TGF-β for 72 hours in U87MG cells. **(F)** Bar graph and immunoblot represents protein expression of DRP1(p=0.0026), pDRP616(p=0.0471), and MFF(p=0.0035) after 72 hours of TGF-β treatment in U87MG cells. GAPDH serves as a loading Control. **(G)** Immunofluorescence study showing DRP1 expression post-TGF-β treatment for 72 hours in U87MG cells. Scale bar: 10µm. Images were analyzed using Zen 2.3 SP1 software. **(H)** Immunoblot analysis representing protein expression of DRP1(p=0.0937) post-TGF-β and N-Acetyl Cysteine treatment was given 2 hours before TGF-β for quenching ROS in U87MG cells**. (I)** The oxygen consumption rate was determined using OCR study in U87MG cells. The results of our study are represented as mean ±SD *(n*=3). A paired, two-tailed t-test was used to determine the significance when two groups were compared. The statistically significant differences are represented with (*), where p<0.05.

### Inhibition of mitochondrial fission restores spatial mitochondrial distribution

An increased mitochondrial fission or elevated expression of DRP1 has often been linked to diverse effects on tumor cells. It can extend from deterioration of mitochondrial health and accompanied apoptosis; to promotion of mitochondrial division whilst mitosis survival of cancer cells by blocking calcium-mediated apoptosis or accentuating the migratory capability of tumorigenic cells [14], [24], [28]. Indeed, a high DRP1 expression was associated with reduced survival of GBM patients as depicted through Kaplan-Meier plot, obtained from human protein atlas and which was confirmed through patient samples analysis that DRP1 expression levels remain high in GBM patients **(Supp. Figure 3A)**. Herein, Mdivi-1 is a pharmacological inhibitor of DRP1 with allied effects like generation of heightened mitochondrial oxidative stress, attenuated mitochondrial metabolism and altered oxygen consumption [29], [30]. Before its application on the GBM cells, we initially analyzed the cytotoxic effect of the compound. Importantly, Mdivi-1 was minimally toxic to the GBM cells, as confirmed through MTT and Annexin V-PI assay. However, there was a distinct alteration in EMT-like associated features in the U87MG cell line **(Supp. Figure 3B & 3C)**. To validate this observation further, we performed a genetic knockdown of DRP1 with siRNA, which showed a similar trend in mitochondrial distribution, just like Mdivi-1. DRP1 inhibition with Mdivi-1 or knockdown with siRNA resulted in a decrease in DRP1 expression levels in U87MG cell line (**Figure 3A, Supp. Figure 3D and 3E)**. Importantly, inhibition of fission using Mdivi-1 or siDRP1 caused a drastically altered localization of the mitochondria from the periphery to the perinuclear region in U87MG cell line (**Figure 3B**). To confirm the findings, we performed staining of the OMM protein-TOM20 in U87MG cell line (**Figure 3C**), and the pharmacological inhibition of fission (Mdivi-1) reversed the localization of TOM20 immunofluorescence as well in U87MG cell line **(Figure 3C)**. This dynamic orientation of mitochondria was further confirmed through live cell imaging studies, as performed through Mito-Tracker Deep Red FM staining establishing the fact that mitochondria indeed undergo dynamic division cycle and movement whilst EMT transformation in U87MG cell line (**Figure 3D**). The change in distribution of mitochondria was confirmed with TOM20, a mitochondrial outer membrane protein and altered expression of DRP1 and their colocalization in U87MG cell line **(Figure 3E**) in. Since three-dimensional studies often more closely replicate a physiological event observed in 2D, we cultured the GBM cells as spheroids. As mentioned earlier, TGF-β-induced EMT was associated with a proliferative arrest of GBM cells. In accordance with the above, we observed a reduced size of spheroids after 3 days of incubation with the cytokine. Interestingly, Mdivi-1 treatment prevented such reduction in spheroid dimensions, reflecting an effect beyond fission, especially on the proliferative status of the U87MG cells as well **(Figure 3F)**. While earlier studies report inhibition of 3D tumor sphere-forming capacity and stemness of breast cancer cells, our observations, in contrast, demonstrates a re-emergence of proliferative potential of Mdivi-1 treated GBM cells thus adding a TGF-β antagonistic function to its wide array of effects. We confirmed through cell cycle analysis that the cells were coming out of the G0-G1 arrest upon Mdivi-1 treatment in U87MG cell line **(Supp. Figure 3F).** These results suggest that mitochondrial fission is an essential prerequisite for mitochondrial localization. Its inhibition leads to the re-distribution of mitochondria.

**Figure 3.**
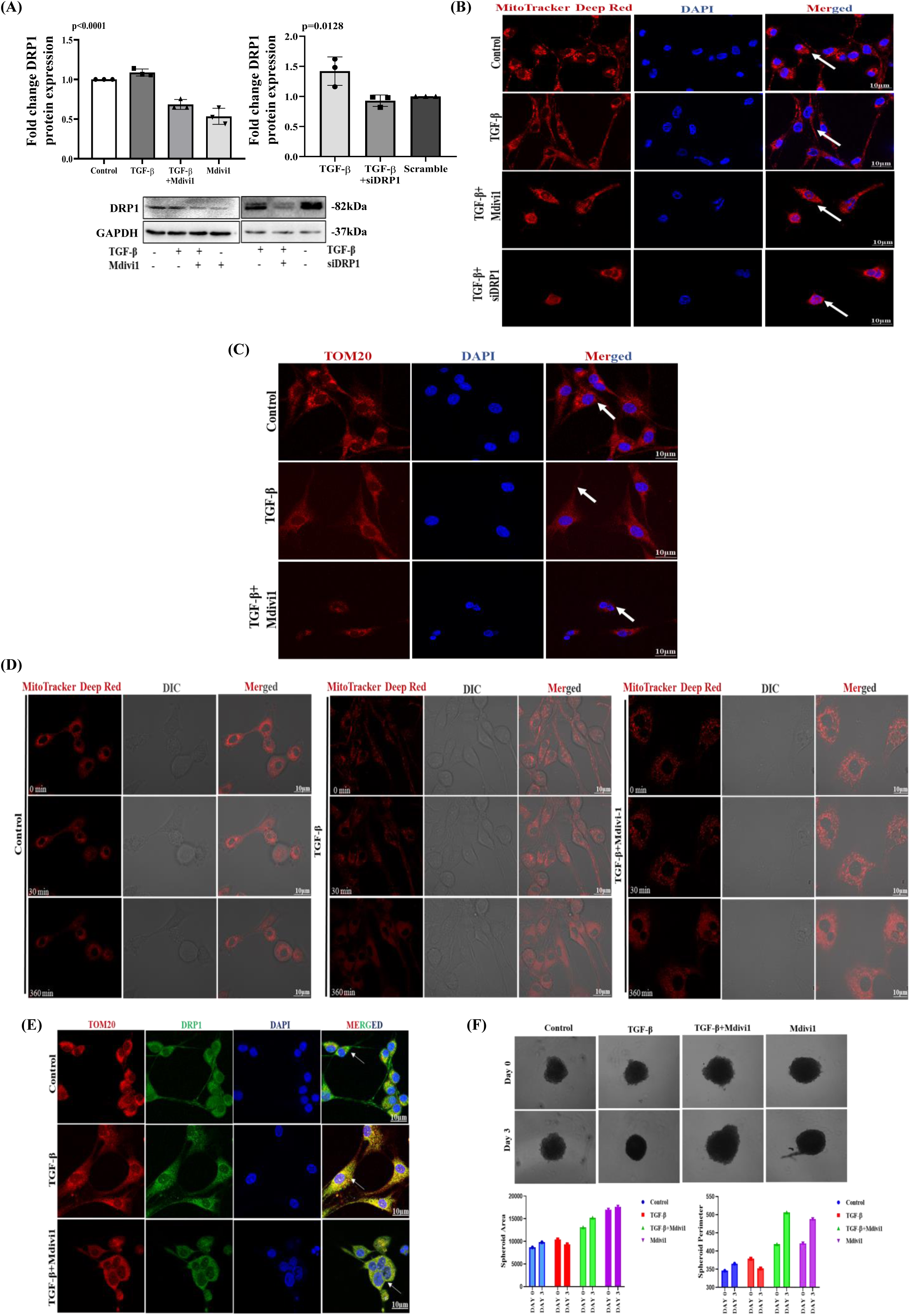
Inhibition of mitochondrial fission restores spatial mitochondrial distribution: **(A)** Bar graph and immunoblot representing DRP1 protein expression post-TGF-β treatment coupled with pharmacological inhibition with Mdivi-1 (10µM) (p<0.0001) or siRNA mediated knockdown of DRP1 (40nM) (p=0.0128) respectively in U87MG cells. **(B)** Mitochondrial staining was done using MitoTracker Deep Red FM post-TGF-β treatment in U87MG cells to study mitochondrial localization upon siRNA-mediated knockdown of DRP1 and via pharmacological inhibition using Mdivi-1 (10µM). Scale bar: 10µm. DAPI (in blue) was used to stain the nucleus. White arrows indicate the localization of mitochondria. **(C)** Immunofluorescence staining shows alterations in TOM20 expression post TGF-β treatment and post-inhibition of DRP1 with Mdivi-1 (10µM) in U87MG cells. **(D)** Live cell imaging using MitoTracker dye shows localization of mitochondria upon TGF-β and Mdivi-1 exposure to control in U87MG cells. **(E)** Immunofluorescence of DRP1 showed differential localization post-TGF-β (20ng/ml) or in combination with Mdivi-1 (10µM) in U87MG cells; DAPI was used to stain the nucleus. Scale bar: 10µm. Microscopic images were captured using Zen 2.3 SP1 software using a 63x oil objective lens under a fluorescent microscope (LSM 880 Axio Observer or Zeiss ApoTome.2). White arrows indicate the localization of TOM20 inside the cell. **(F)** U87MG cells were cultured in 3D spheroids, and phase contrast images were taken using Zen2.3 SP1 software using 10x objective; Scale bar 100μm. Bar graphs represent a change in the spheroid area and perimeter, post-TGF-β (20ng/ml) or Mdivi-1 (10µM) exposure for about 3 days.

### Inhibition of fission causes EMT reversal or DRP1 knockdown leads to reduced Glioma cell invasion and F-actin remodeling

Apart from the reversal of growth inhibitory effects of under TGF-β the other striking feature observed in the Mdivi-1 or siRNA-treated cells was a change in cell shape. To confirm the same, we performed SEM imaging of U87MG cell line, and as mentioned above, careful observation of the cells revealed that upon DRP1 inhibition, cells significantly altered their morphology and obtained a considerably rounder phenotype (**Figure 4A**). This prompted us to check for the association of mitochondrial fission with EMT. We hypothesized that mitochondrial dynamicity and localization dictate TGF-β-induced EMT. Interestingly, the transcriptional levels of EMT markers in U87MG cell line were significantly downregulated upon inhibition of mitochondrial fission (**Figure 4B**). In parallel, a significant reduction in protein levels of EMT markers-Vimentin and N-cad was also obtained upon treatment with Mdivi-1 in GBM cell lines (**Figure 4C and Supp. Figure 3E**). There was also a reduction in the immunofluorescence intensity of Vimentin in U87MG cell line (**Figure 4D)**. Importantly, si-RNA-mediated knockdown of DRP1 also resulted in a similar reversal of EMT features in U87MG cell line (**Figure 4E)**. The above results indicate that mitochondrial dynamicity can potentially regulate the differentiation of GBM cells. Herein, several researchers have also demonstrated that actin polymerization at or near fission sites can regulate DRP1’s activity [31], [32]; it can dynamically bind to actin filaments and aid in oligomeric maturation of DRP1 [33], [34]. Interestingly, in our study, phalloidin staining showed the altered distribution of actin fibers, more to the perinuclear region post-Mdivi-1 or siDRP1 treatment compared to TGF-β suggesting an F-actin re-organization as well, upon inhibition of fission in U87MG cell line (**Figure 4F**). Finally, when the glioblastoma cells were given a Mdivi-1 rescue, in presence of TGF-β, it resulted in a re-emergence of EMT phenotype thus reflecting that the differentiation event orchestrated by cytoskeletal and mitochondrial distribution is dynamic in U87MG cell line (**Figure 4G**).

**Figure 4.**
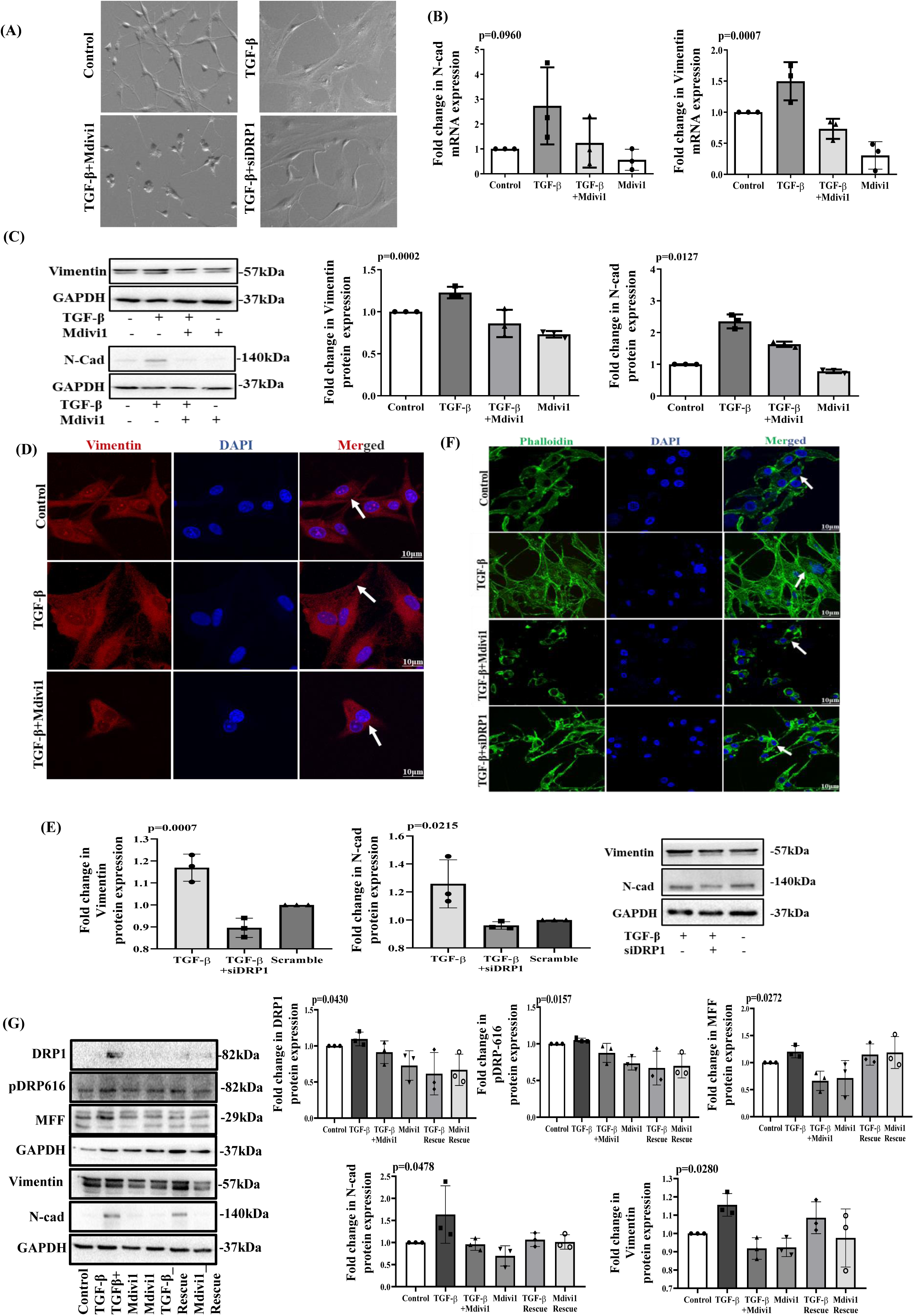
Inhibition of fission causes EMT reversal: **(A)** Scanning electron microscope (SEM) images of U87MG cells after treatment with TGF-β (20ng/ml) alone or in combination with Mdivi-1 (10µM) or siDRP1 (40nM) for 72 hours; Scale bar 100μm. White arrows indicate a change in cellular morphology. Scale Bar: 10µm **(B)** Bar graph representing a change in mRNA expression of N-cad (p=0.0960) and Vimentin (p=0.0007) post-TGF-β (20ng/ml) treatment for 72 hours in 87MG cell line. **(C)** Bar graphs and immunoblot representing EMT markers Vimentin (p=0.0002) and N-cad (p=0.0127) protein expression post-TGF-β (20ng/ml) and in combination with Mdivi-1 (10µM) for 72 hours in U87MG cell line. GAPDH serves as a loading control. **(D**) Immunofluorescence images of Vimentin in U87MG cell line. White arrows indicate the difference in protein localization upon TGF-β (20ng/ml) exposure, alone or combined with Mdivi-1 (10µM), Scale bar: 10µm. **(E)** Bar graphs and immunoblots represent change in Vimentin(p=0.0007) and N-cad(p=0.0215) protein expression in U87MG cell line post DRP1-knockdown using siRNA (40nM) for 72 hours. GAPDH serves as a loading control. **(F)** F actin fibers in U87MG cells were stained with phalloidin (Green), and the nucleus was stained with DAPI (blue). Scale bar: 10µm. White arrows indicate the spread of F-actin filaments in the cell. **(G)** Bar graphs and immunoblot shows DRP1(p=0.0430), pDRP616(p=0.0157), MFF(p=0.0272), Vimentin(p=0.0478), and N-cad(p=0.0280) protein expression in U87MG cells post-Mdivi-1 (10µM) rescue for 72 hours. When comparing two groups, the significance was ascertained using paired, two-tailed t-tests and one-way ANOVA with Bonferroni’s test for multiple comparisons. The following p values relate to the statistically significant differences from the control group: *p ≤ 0.05, **p ≤ 0.01, ***p ≤ 0.001. The considerable variations are indicated with (*).

### RhoA regulates DRP-1 mediated mitochondrial fission in GBM cells

The above results indicate that the cytoskeleton and mitochondria have intense crosstalk governing cellular differentiation events like EMT. Herein, we questioned whether modulation of the cytoskeleton or actin organization induced under TGF-β can govern the cytokine-induced mitochondrial fission. Interestingly, inhibition of actin polymerization using the drug Latrunculin inhibiting myosin activity using Blebbistatin or Dynamin inhibitor-dynasore resulted in a withdrawal of EMT features as observed through SEM validating the link between actin regulation and EMT in U87MG cell line **(Figure 5A).** It also induced peri-nuclear localization of the mitochondria in U87MG cell line **(Figure 5B).** Importantly, the inhibition of actin polymerization and myosin activity led to downregulation of EMT markers, along with mitochondrial fission proteins like DRP1 and MFF confirming the hypothesis that actin regulation can also dictate mitochondrial fate-fission/fusion in the GBM cells in presence of TGF-β in U87MG cell line **(Figure 5C).** Herein, a protein known to play a crucial role in actin assembly is RhoA; it can not only drive structural alterations involved in cytoskeleton re-organization but can also coordinate other cellular events like, transcriptional activation, tumor cell proliferation, survival, migration and invasion [35], [36]. To validate the same, we initially analyzed the expression of RhoA upon EMT induction, and it showed a drastic increase, suggesting that a connection might exist between RhoA and molecules regulating EMT in U87MG cell line **(Figure 5D)**. Interestingly, a co-immunoprecipitation analysis revealed that RhoA physically interacts with DRP1. Therefore, we hypothesized that this interaction might be crucial for mitochondrial structure, distribution, and EMT induction, as observed with TGF-β exposure in U87MG cell line **(Figure 5E).** Importantly, inhibition of RhoA activity with Y27632 resulted in an attenuation in expression of both EMT markers and mitochondrial fission associated proteins as well in U87MG cell line **(Figure 5F)**. The above findings thus demonstrate that TGF-β induced RhoA plays a key role in not only coordinating actin re-organization and cytoskeletal modifications conducive to EMT but also participates in regulation of mitochondrial fission that is necessary for the EMT process.

**Figure 5.**
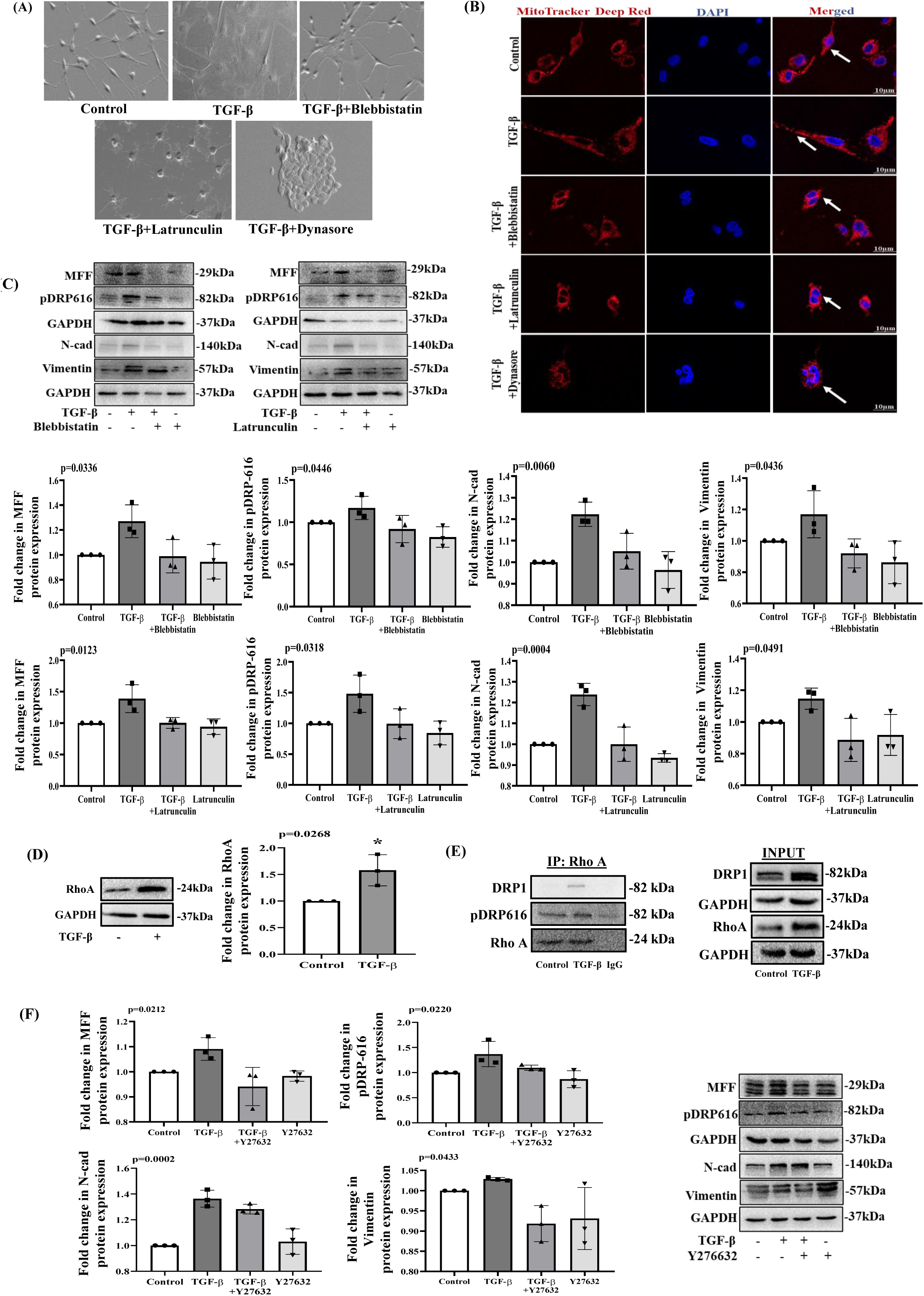
RhoA regulates DRP-1 mediated mitochondrial fission in GBM cells: **(A)** Scanning electron microscope (SEM) images of U87MG cells after treatment with TGF-β (20ng/ml), either alone or in combination with Blebbistatin (10µM), Latrunculin (100ng/ml) and Dynasore (80µM) respectively. The white arrows indicate changes in cellular morphology post the respective treatment regime; Scale bar 100μm. **(B)** Mitochondrial localization was represented using MitoTracker Deep Red FM staining post-exposure with TGF-β (20ng/ml) and prior-mentioned combinations in U87MG cells.; Scale bar: 10µm. White arrows indicate changes in mitochondrial localization upon different drug treatment regimes. **(C)** Changes in MFF(Blebbistatin; p=0.0336 & Latrunculin; p=0.0213), pDRP616(Blebbistatin; p=0.0446 & Latrunculin; p=0.0318), N-cad(Blebbistatin; p=0.0060 & Latrunculin; p=0.0004), and Vimentin(Blebbistatin; p=0.0436 & Latrunculin; p=0.0491), protein expression through Bar graph and immunoblot, post-exposure with TGF-β (20ng/ml), and combination with Blebbistatin (10µM) and Latrunculin (100ng/ml) respectively in U87MG cell line. (**D)** Immunoblot and bar graph analysis represent the expression of RhoA (p=0.0268) post-TGF-β (20ng/ml) exposure in U87MG cell line. GAPDH served as a loading control. **(E)** Co immunoprecipitation with RhoA antibody, followed by immunoblotting for DRP1, pDRP616, and RhoA in both immuno precipitated and total cell lysate (input) samples (*n* = 3) in U87MG cell line. **(F)** Changes in MFF(p=0.0212), p-DRP616(p=0.0220), N-cad(p=0.0002), and Vimentin(p=0.0433) protein expression levels upon exposure with TGF-β, as well as in combination with a pharmacological inhibitor of RhoA: Y27632(25µM) in U87MG cell line. Unless otherwise mentioned, a paired t-test was used for statistical analysis, where p<0.05 was considered significant and represented with (*).

## 1. Discussion

Mitochondria play a vital role in multiple cellular processes, including cellular energy production, oxidative stress, and overall metabolic homeostasis. Due to their altered metabolism, high energy needs, and elevated ROS tolerance, we hypothesized that mitochondria play a central role in cancer pathogenesis[13], [15] In this context, we investigated mitochondria’s contribution, especially in one of the critical events associated with cancer cell invasion-epithelial-mesenchymal transition (EMT). The objective was to understand the key interactions between mitochondria and EMT, which can provide valuable insights into potential targets for preventing subsequent cancer metastasis [11] In this regard, the model of our study was glioblastoma, a highly aggressive tumor often showing intracranial metastasis involving EMT-like transitions, which can also contribute to resistance to conventional therapies. Characterization of EMT markers in the presence of TGF-β, a cytokine highly prevalent in the GBM micro-environment, showed robust EMT phenotype in the GBM cells studied. In parallel, analysis of patient samples also indicated increased expression of EMT-markers and its association with poor prognosis in GBM. Given the existing challenge in GBM therapy, we focused on identifying key factors, such as mitochondrial dynamics, that regulate this critical step of the metastatic cascade. Interestingly, the number of mitochondria in GBM cells transiently increased, accompanied by a high mitochondrial ROS and an up-regulated mitochondrial fission, whilst cells manifested EMT in response to TGF-β. A fragmented mitochondrial morphology was coupled to the up-regulation of fission markers like MFF, DRP1, and FIS1. It has been discovered recently that several human malignancies exhibit up-regulated DRP1 expression [14], [24], [28]. Therefore, we gained a more profound comprehension of DRP1 expression in gliomas, which showed that a higher protein expression is associated with comparatively poor survival. Our patient data analysis also indicated that gliomas have a higher expression of the fission mediator-DRP1. To further validate, we demonstrated that down-regulation of DRP1 expression in glioma reduced EMT *in vitro* with a noticeable shift in the actin cytoskeleton. In this regard, the RhoA-Rho kinase (ROCK) pathway has been prominent in recent times as a regulatory mechanism of the actin cytoskeleton [35].

Furthermore, prior research has demonstrated that RhoA downregulation can reduce glioma cell activities such as invasion, migration, and proliferation via modifying actin cytoskeleton remodeling [36]. Interestingly, in our investigation, actin polymerization inhibition reduced RhoA levels and consequently caused a downregulation of mitochondrial fission. We accordingly hypothesized an interaction between DRP1 and RhoA signaling pathway, which was confirmed through a co-immunoprecipitation study. We provide conclusive evidence for regulating GBM invasiveness through crosstalk between RhoA signaling and DRP1 controlling cytoskeleton remodeling. As a dynamic cellular organelle, mitochondria maintain equilibrium in terms of number and morphology, as well as through mitochondrial quality control and turnover. Though earlier studies on renal cells have correlated TGF-β activity with mitochondrial dysfunction, till date, there are limited reports highlighting the direct role of fission-associated proteins and their regulation by master cytoskeletal regulators like RhoA dictating EMT and aggressive features of GBM. Gliomas have historically been clinically challenging due to their infiltrative growth patterns and high rate of malignant recurrence, and therefore molecular insights identifying targetable markers mitigating the above conditions can be an effective strategy addressing treatment bottleneck. However, further studies delineating the role of mitochondria, RhoA, and their targeting in vivo are required to establish novel future therapeutic modalities.

## Materials and Methods

### Cell culture

A human glioblastoma (U87) cell line was obtained from the National Centre for Cell Science (NCCS) in Pune. A minimal essential medium containing Earle’s salt (EMEM; Hi-media, AL047S) supplemented with 10% fetal bovine serum (FBS; Invitrogen, 26140-079) and antibiotics (1% penicillin-streptomycin solution; Invitrogen, #10378-016) was used to grow the cells at 37°C and 5% CO_2_. Glioblastoma cells were grown to 70% confluency prior to cytokine treatment. They were then aspirated, rinsed with phosphate-buffered saline (PBS), and grown for 12 hours in a serum-starved medium (EMEM with 1% FBS). For 72 hours, TGF-β (20 ng/ml) was the cytokine employed for therapy[37].

### Cell viability assay

We performed the MTT assay to evaluate cell viability. B riefly, U87 cells were grown in 96-well plates, serum-starved for 12 hours, and then exposed to varying cytokine or drug doses. After adding 20 µl of MTT (stock concentration: 5 mg/ml), the cells were incubated for four hours. After dissolving formazan crystals in DMSO, measurements were taken at 570 nm using a 630 nm differential filter using a Multiskan Sky spectrophotometer (Thermo Scientific, 90273020) to assess viable cells. The treated sample’s percentage viability was compared to the control’s. The formula viability (%) = (mean absorbance value of drug-treated cells)/(mean absorbance value of control) * 100 was used to determine the proportion of viable cells[38].

### RNA isolation, cDNA Synthesis, and Quantitative Analysis using RT-PCR

Glioblastoma cells were treated with a cytokine or inhibitory drugs and allowed to grow in 10 cm dishes. TRIzol reagent (Sigma, T9424) was used to extract post-treatment whole-cell RNA, which was then measured and its purity checked. RNA treated with DNase I produced first-strand complementary DNA (cDNA) (Thermo Scientific, EN0521). An iScript cDNA synthesis kit (Bio-Rad, #1708891) with oligodT was used to create cDNA in accordance with the manufacturer’s instructions. Gene-specific primer sequences were used to amplify the templates (see Table 1 below). To detect B-actin or GAPDH, SYBR Green Supermix (Bio-Rad, #170-8882AP) were used as a housekeeping control[37]. Pfaffl’s technique was used to calculate the relative mRNA expression. For Vimentin, N-Cad, MFF, DRP1 & FIS1, and GAPDH the melting temperature for the PCRs were 53 °C, 55 °C, 63.3 °C, 57.2 °C, and 62.1 °C, respectively.

**Table.1.**
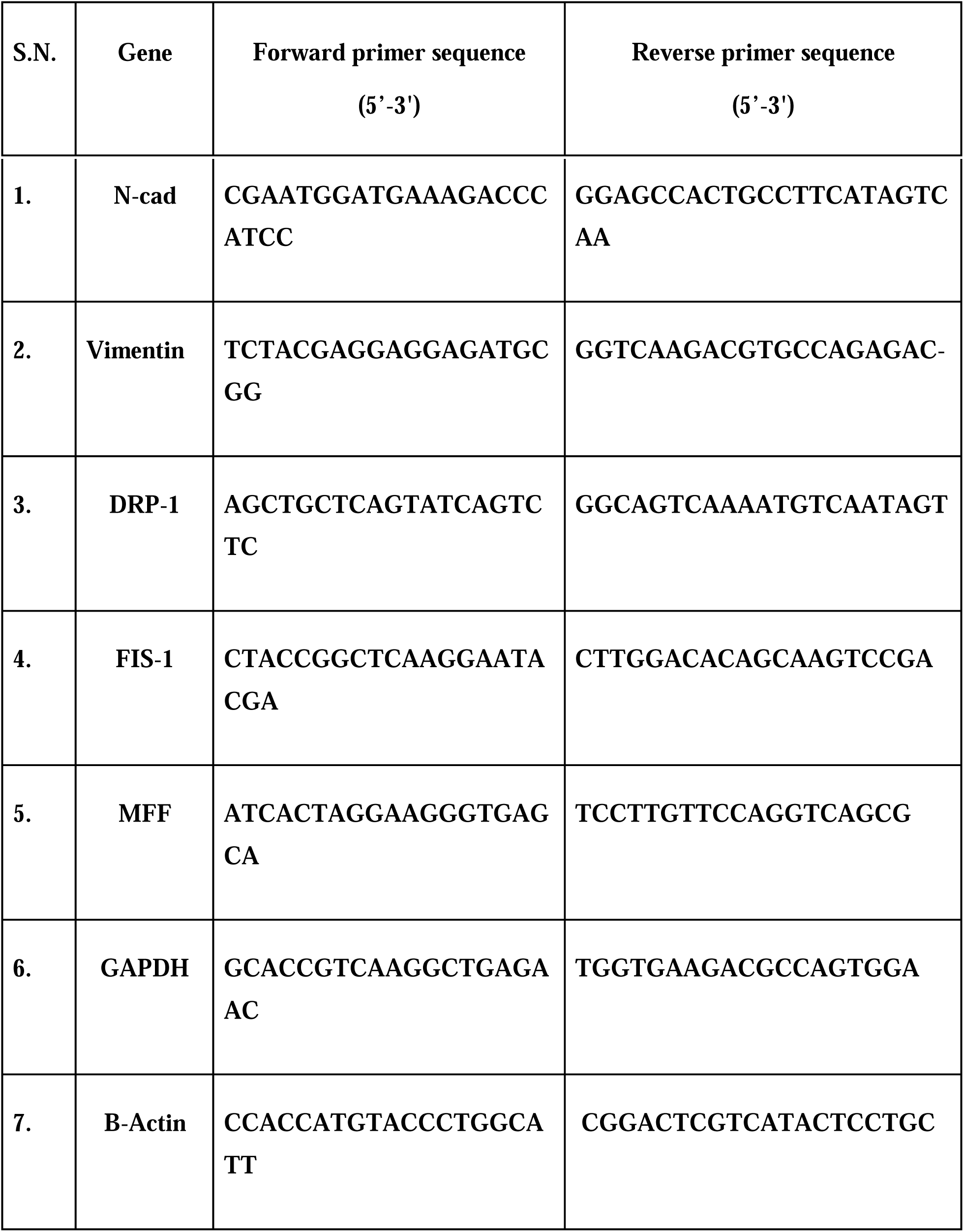

### Transfection

Cells were grown upto 70% density in each well of the six-well plates. Following the manufacturer’s recommendations, lipofectamine 3000 reagents were used to transfect the cells with a particular siRNA (40nM or 60nM)[39].

### Co-immunoprecipitation and Immunoblotting

RIPA lysis buffer (#R0278, Sigma-Aldrich) with protease inhibitor (#P8340, Sigma-Aldrich) was used to extract proteins from U87 cells. Bradford reagent (Sigma) was used to estimate the total protein concentration at a wavelength of 595 nm. The antibody for RhoA and DRP1 (1 μg/uL) was used to pull down 500 μg of protein in total. After being run on denaturing polyacrylamide gels, the cellular protein lysates were transferred to a polyvinylidene fluoride membrane (PVDF, BioRad) and blocked with 5% skim milk (HiMedia). After that, specific primary antibodies (at dilution 1:1,000) were used to probe and re-probe the blots. Following the manufacturer’s instructions, the secondary antibodies (at dilution 1:2,000) were conjugated with horseradish peroxidase and identified using an ECL detection system with ChemiDoc (Bio-Rad)[37]. The blots were then cut to probe with different antibodies against proteins of varying molecular weights where required. The ImageJ program was used to quantify the expression of different proteins, and GraphPad Prism 8.0.1 was used for analysis.

### Immunofluorescence microscopy

After being grown on coverslips, cells were treated with cytokine and other drugs once the required confluency was achieved. The media was then taken out, and PBS wash was given. For 15 minutes, cells were fixed at room temperature using 4% paraformaldehyde. Following the PBS wash, cells were permeabilized for two minutes at room temperature using Triton-X 100 (0.2%). Additionally, cells were blocked with 2.5% bovine serum albumin (BSA) for 1 hour at room temperature followed by PBS washes. After being incubated with the primary antibody for overnight at 4°C, the cells were rinsed three times with PBS before being incubated for an hour with the Alexa flor 488 or Alexa flor 555 conjugated secondary antibody[40]. Slides were prepared using antifade DAPI, and several PBS washes were given. A fluorescent microscope (LSM 880 Axio Observer, or Axio Observer.Z1/7) was used to observe the slides. Zen 3.2 (blue edition) software was used for analyzing microscopic images.

### MitoTracker Deep Red (MTR) staining

MitoTracker Deep Red FM was used to stain the mitochondria. Following treatment, the cells were incubated at 37°C for 20 minutes with MitoTracker Deep Red FM. The coverslips were washed with PBS, fixed with 4% paraformaldehyde at room temperature, and then mounted with antifade DAPI. To identify the position and shape of the mitochondrial pictures, the cells were looked at using a ZEISS Axio Scope A1 microscope with ZEN 3.2 (blue edition). Image J software was then used for analysis[40].

### Cellular ROS assay

To estimate Reactive oxygen species inside cells, w e used 2,7-dichlorofluorescein diacetate (H2DCF-DA) (Sigma) which assess the amounts of ROS inside cells. Before administering any treatment, cells were seeded in 96-well plates at a density of 8 × 10^3^ cells/well and exposed to the ROS scavenger NAC (10 mM) for two hours. In the ROS studies linked to DRP-1 inhibition, Mdivi-1 was introduced after ROS were quenched with NAC and then TGF-β. After cytokine or drug exposure, the cells were rinsed with PBS and incubated with DCFDA (10 mM) for about an hour[23]. A microplate reader was used to quantify fluorescence at 530 nm emission and 485 nm excitation (Fluoroskan Ascent). Using CytExpert Software, samples were passed through a flow cytometer to measure the fluorescence intensity.

### JC-1 assay

The JC-1 assay was employed to evaluate the mitochondrial membrane’s potential upon TGF-beta treatment. TGF-β was administered for 72 hours after the cells were seeded at a density of 60,000 cells/well in a 6-well plate, allowed to adhere, and then serum starved with 1%FBS containing EMEM medium. After treatment, the cells were incubated for 15 to 30 minutes at 37 oC with 5% CO_2_ in the dark with 75 nM JC-1 dye (working: 1 μl JC-1 from stock in 12 mL of complete EMEM). Post treatment, the cells were taken out, re-suspended in 1X PBS, and then sent directly for analysis using flow cytometry (Cytoflex, Beckmann Coulter)[40]. CytExpert software was used to analyze the data obtained throughout this process.

### Estimation of mitochondrial ROS generation

The amount of Superoxide generated by the mitochondria of particular cells was measured using MitoSOX Red, which was acquired from Invitrogen. In a 6-well plate, the GBM cells were seeded. To assess the amount of superoxide generated by the mitochondria, after attaining their morphology and confluency, cells were first serum deprived with 1% FBS containing EMEM media and then treated with TGF-β for 72 hours, respectively. Following the cytokine exposure, each well was incubated with 1 mL of 5 μM MitoSOX and was left at room temperature for ten minutes. The cells were then given a PBS wash. The cells were immediately transferred for flow cytometry analysis after being extracted and re-suspended in 1X PBS (Cytoflex, Beckmann Coulter)[40]. CytExpert software was used to analyze the data gathered throughout this process.

### Phalloidin staining

Cells were treated with cytokines, rinsed with PBS, and then fixed in 2% paraformaldehyde for 10 minutes at room temperature in the dark. Cells were subjected to a 0.2% Triton X 100 solution for 2 minutes at room temperature followed by several PBS washes. Further, the cells were incubated for approximately 30 minutes at room temperature in the dark with phalloidin stain (1:2000 in 1% BSA). The cells over coverslip were stained with DAPI and after several PBS washes, the slides were prepared. Slides were further examined under a fluorescent confocal microscope (LSM 880 Axio Observer)[38].

### Scanning electron microscope (SEM)

After seeding cells on a coverslip, cytokine treatment was given. The media was disposed of after the cytokine incubation, and PBS wash was administered. After that, cells were fixed with 4% formaldehyde and left at room temperature for ten minutes in the dark. After many PBS washes, cells were placed in ethanol in gradients of 25%, 50%, 75%, and 100% for one minute each. The cells were allowed to air dry after being dipped in 100% ethanol. The coverslips were coated with conductive gold, and a SEM (Thermo Fisher/Fei; Apreo S) was used for microscopy[38].

### Annexin/Propidium iodide (PI) staining

In a six-well plate, cells were seeded and they were allowed grow until 70% confluency. The cells were collected after being exposed to cytokines or drugs, rinsed with PBS, and then re-suspended in 500 μl of 1X Annexin binding buffer. After adding Annexin V/PI, the cells were placed in the dark for 20 minutes. To estimate the proportion of apoptotic cells, they were immediately run through flow cytometer (Cytoflex, Beckmann Coulter). CytExpert software was used to evaluate the collected data. Apoptotic cells were quantified by counting the proportion of cells in the lower right and upper right quadrants (LR-early apoptotic and UR-late apoptotic, respectively)[23].

### Cell cycle analysis

Cells were grown in a six-well plate in order to analyze the DNA content. Following the cytokine or drug treatment, the cells were collected in an Eppendorf, rinsed with PBS, and centrifuged for 5 minutes at 2000 rpm and 4° C to pellet them down. 100 µl of ice-cold PBS was used to re-suspend the pellet, and 900 µl of ice-cold 70% ethanol was added dropwise while being continuously stirred. For fixing, the cells were stored at -20°C for overnight. The next day, pellet was re-suspended in 450 µl of PBS with 10 µl of propidium iodide (PI; 2 mg/ml) after the cells were centrifuged (21). Followed by 10 minutes of incubation in dark, a flow cytometer (Cytoflex, Beckmann Coulter) was used for analysis, and CytExpert software was used to examine the results[38].

### Patient sample RT-PCR

Surgically resected GBM tumor tissues were collected from the Department of Neurosurgery, AIIMS, Delhi, after due consent from patients and ethical clearance from the institute ethics committee (<u>IEC-711/07.08.2020, RP-43/2020</u>). Tumor tissues were collected in RNA-Later solution (Ambion, USA, catalog no. AM7020), incubated at 4°C for 24 hours, and transferred to -70°C after removal of RNA-Later solution until further use. The histopathological diagnosis of tumors was done by neuropathologist Prof. Chitra Sarkar/ Prof. Vaishali Suri. Total RNA from frozen tumor samples was extracted using TRI Reagent (Sigma-Aldrich, USA) and quantified using NanoDrop ND-1000 spectrophotometer (Thermo Fisher Scientific, USA). Genomic DNA was removed by DNase I (MBI Fermentas, Hanover, MD) treatment [41]. The reverse transcriptase enzyme (MBI Fermentas, Hanover, MD) and random decamers (MWG, India) were used in 20μl reaction volumes to conduct reverse transcriptase reactions. The control for the q-PCR analysis was normal human brain total-RNA (Clontech, USA, catalog number 636530). NanoDrop was used to measure the RNA’s quality and concentration. Each sample included 1000 ng of RNA for cDNA synthesis. Following the manufacturer’s instructions, random hexamer was employed in the Verso cDNA synthesis kit (Thermo Fisher Scientific, K1632). For cDNA synthesis, a total of 20µl reactions were prepared. Following cDNA synthesis, 2.5 µl of the reaction was used for each qPCR experiment after it had been diluted to a 1:5 ratio in nuclease-free water. The SYBR™ Green PCR Master Mix (Thermo Fisher Scientific, 4309155) was used to conduct SYBR green-based qPCR for transcript level quantification. The qPCR involved 5 minutes of initial denaturation at 95°C, which was followed by 35 cycles of 10 seconds at 95°C, 45 seconds at 60°C, and 30 seconds at 72°C, and a final extension of 10 minutes at 72°C. The 2−ΔΔCT-method was used to compare the relative expression of the examined genes to that of the housekeeping gene 18s. For every condition, three separate experiments were carried out in triplicate.

### Determination of oxygen consumption rate

Cells were seeded in a 96-well plate at a density of 6000 cells per well, and incubated overnight in a CO_2_ incubator. Cytokine treatment was given, as discussed above. Once the treatment is over, replenish the old media with fresh media. Added extracellular O_2_ consumption reagent and promptly sealed each well with pre-warmed high-sensitivity mineral oil per the manufacturer’s protocol. Readings were taken in a fluorescence plate reader to measure the O_2_ consumption signal at 1.5-minute intervals for 90-120min and normalized the readings with the cell viability.

### In-silico analysis

We conducted a comprehensive in silico analysis using the GSE50161 gene expression dataset from the NCBI-Gene Expression Omnibus (GEO) repository (https://www.ncbi.nlm.nih.gov/geo/query/acc.cgi?acc=GSE50161). The dataset, comprising profiles from human brain tumors and normal brain tissues, was analyzed using R to identify differentially expressed genes (DEGs) through statistical methods. The list of dysregulated genes was subjected to Gene Ontology (GO) enrichment analysis via the ShinyGO 0.80 web tool (http://bioinformatics.sdstate.edu/go/). This tool facilitated the categorization of the dysregulated genes into biological processes, cellular components, and molecular functions. The enriched pathways were visualized using bar charts highlighting key cellular mechanisms, including immune response, cell cycle regulation, and signal transduction. This computational approach provided insights into the molecular alterations in brain tumor biology.

### Statistical analysis

GraphPad Prism software, version 5.0, was used to analyze the data. One-way or two-way ANOVA was used to statistically assess the outcome of a particular treatment in comparison to the control. The Bonferroni method was used to examine multiple comparisons. The difference was regarded as not significant (ns) if the p-value was greater than 0.05; however, it was considered significant and represented by the symbols * if the p-value was less than 0.05, ** if it was less than 0.01, and *** if it was less than 0.001.

## Supplementary Figures

**Supplementary Figure. 1**. **(A)** Differential expression of biological processes was determined using gene ontology analysis using Shiny GO v0.61. **(B)** Bar graphs and immunoblot show the change in N-cad(p=0.0064, p=0.1920) and Vimentin (p=0.0563, p=0.0200) protein expression in different GBM cell lines: U373 and LN229. B-Actin served as a loading control. **(C)** Bar graph representing dose and time kinetics study of TGF-β in U373, LN229, and U87MG cell lines, respectively. **(D)** Flow cytometric analysis shows cells at various cell cycle stages post TGF-β in U373, LN229 and U87MG cell lines, respectively. **(E)** Bar graph and immunoblot representing the change in p21 (p=0.0003) and PCNA (p=0.0038) protein expression post-TGF-β (20ng/ml) exposure for 72 hours in U87MG cell line. GAPDH served as a loading control. **(F)** Cell viability (MTT assay) post-treatment different concentrations of SIS3 (p<0.0001) and SB431542 (P=0.0169), 10µM of both was used as the inhibitory dose for 72 hours in U87MG cell line. When comparing two groups, the significance was ascertained using paired, two-tailed t-tests and one-way ANOVA with Bonferroni’s test for multiple comparisons. The following p values relate to the statistically significant differences from the control group: *p ≤ 0.05, **p ≤ 0.01, ***p ≤ 0.001. The considerable variations are indicated with (*).

**Supplementary Figure 2. (A)** Flow cytometric analysis using Mito-SOX red and bar graph showing mtROS shift post-TGF-β (20ng/ml) exposure for 72 hours in the U373 Cell line(p=0.0464). **(B)** Analysis of intracellular ROS levels (p=0.1210) and associated cell viability (p=0.07890 following 72-hours exposure to TGF-β in the U87MG cell line. **(C)** Assessment of the alterations in mitochondrial morphology following a 72-hour exposure to TGF-β (20ng/ml) in U373 and LN229 cell lines. MitoTracker Deep Red staining was used to stain mitochondria, and DAPI (blue) stained the nucleus in both cell lines and bar graphs represents the fold change in mitochondrial circularity (p≤0.0001) and fragmentation (p<0.0001). Image-J software was used to assess mitochondrial fragmentation. Scale bar: 10 µm. When comparing two groups, the significance was ascertained using paired, two-tailed t-tests and one-way ANOVA with Bonferroni’s test for multiple comparisons. The following p values relate to the statistically significant differences from the control group: *p ≤ 0.05, **p ≤ 0.01, ***p ≤ 0.001. The considerable variations are indicated with (*).

**Supplementary Figure 3. (A)** Kaplan-Meier plot obtained from human protein atlas represents the correlation between DRP1 (DNML1) expression and survival of GBM patients. The blue and pink curves represent low and high protein expression, respectively, and the bar graph represents mRNA expression levels of DRP1 in normal brain tissue vs GBM patient RNA samples (p=0.2633). **(B)** Analysis of cell viability in Mdivi-1 (10 μM) and after knockdown with siDRP1(40nM) treated cells using Annexin V-PI staining(p<0.0001) in U87MG cell line. **(C)** Analysis of cell viability in Mdivi-1 (dose kinetics) treated cells using MTT assay(p<0.0001) in U87MG cell line. **(D)** Changes in mRNA levels of DRP1 post-exposure to TGF-β (20ng/ml), either alone or in combination with Mdivi-1 (10µM) (p=0.1330) or siRNA (40nM) mediated knockdown of DRP1 (p=0.0199) respectively in U87MG cell line. **(E)** Bar graph and immunoblot representing DRP1(U373;p=0.0081 & LN229; p=0.0004), N-cad(U373;p=0.0155 & LN229; p=0.0245), and Vimentin(U373;p=0.0452 & LN229; p=0.0147) protein levels upon TGF-β (20ng/ml) and in combination with Mdivi-1 (10µM) in U373 and LN229 cell lines. **(F)** Flow cytometric study shows cells at various stages of the cell cycle post-TGF-β (20ng/ml), Mdivi-1 (10µM) alone or in combination in the U87MG cell line. When comparing two groups, the significance was ascertained using paired, two-tailed t-tests and one-way ANOVA with Bonferroni’s test for multiple comparisons. The following p values relate to the statistically significant differences from the control group: *p ≤ 0.05, **p ≤ 0.01, ***p ≤ 0.001. The considerable variations are indicated with (*).

## Data availability

Data will be made available on request.

## Acknowledgments

The authors acknowledge BITS Pilani, Pilani campus, for providing them with infrastructural support. The authors acknowledge Dr. Chitra Sarker and Dr. Vaishali for their help in the tumor diagnosis.

## Authors contributions

RC and SM acquired funding and managed the project. MC^1^ performed the *in-vitro* experiments and analyzed the data. MC^2^, BB, NM, and KC collected the patient sample, performed the RT-PCR, and analyzed the data. SK and SC did the *in-silico* analysis. MC^1^, RC, and SM contemplated the idea and wrote the manuscript. MC^1^, BB, RC, and SM edited the final manuscript.

