## Supplementary Figures for "DRP1-mediated mitochondrial dynamics orchestrate EMT in glioblastoma cells"

**Supplementary Figure 1(A)**

### Biological processes

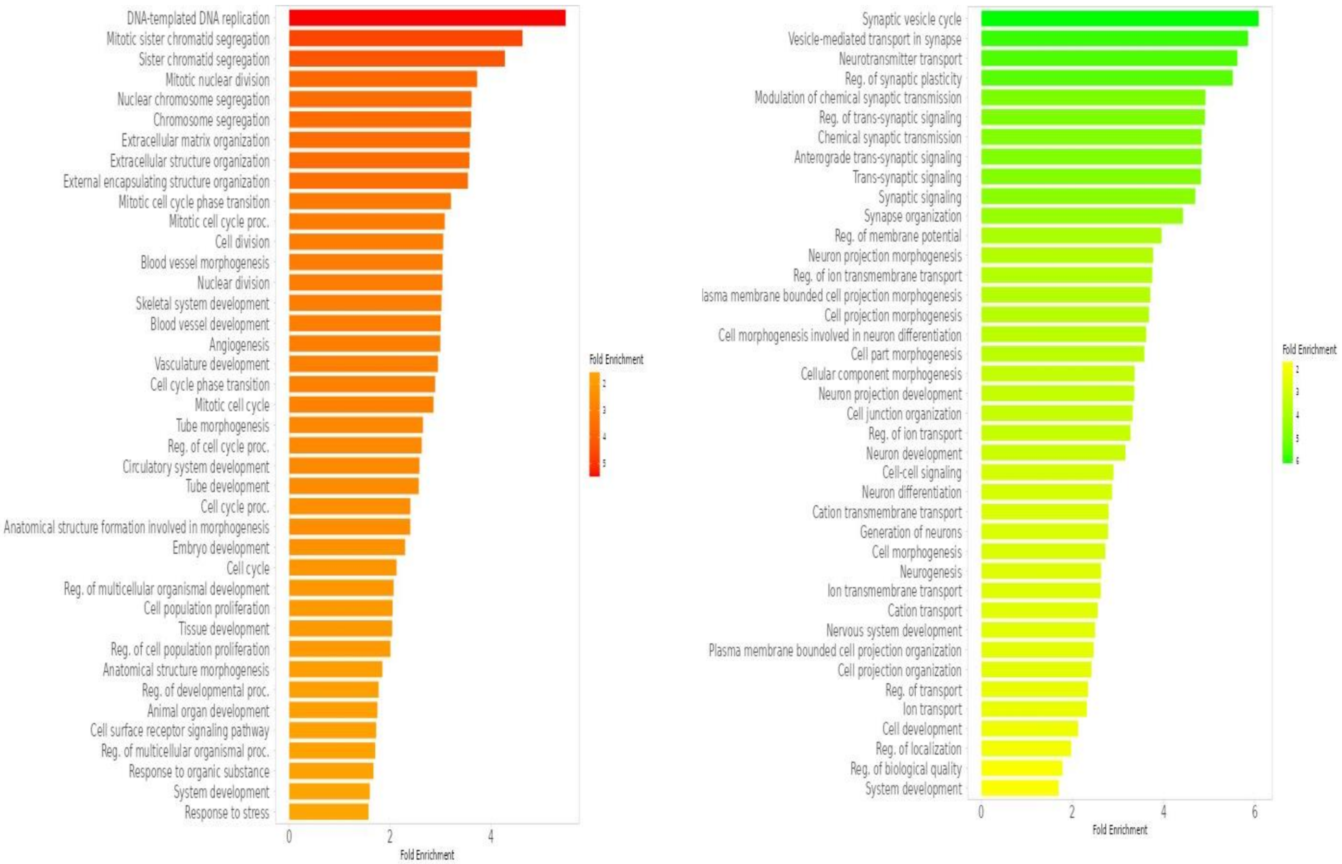

**Supplementary Figure 1(B-E)**

**(B)**

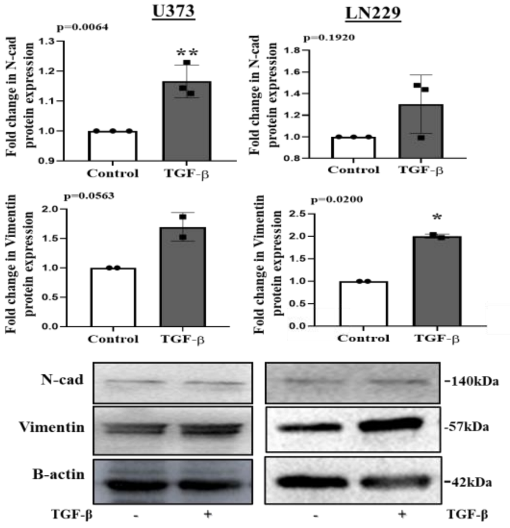


**(C)**

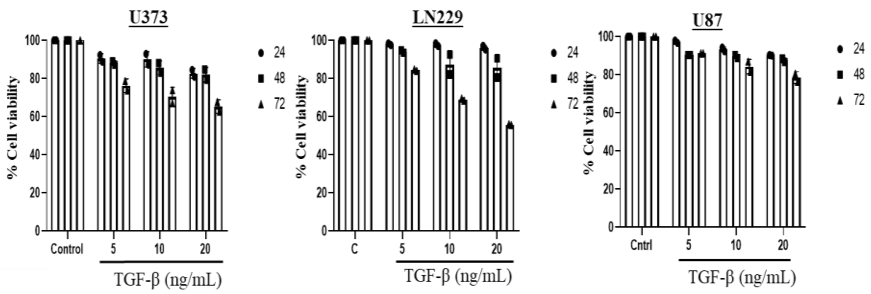


**(D)**

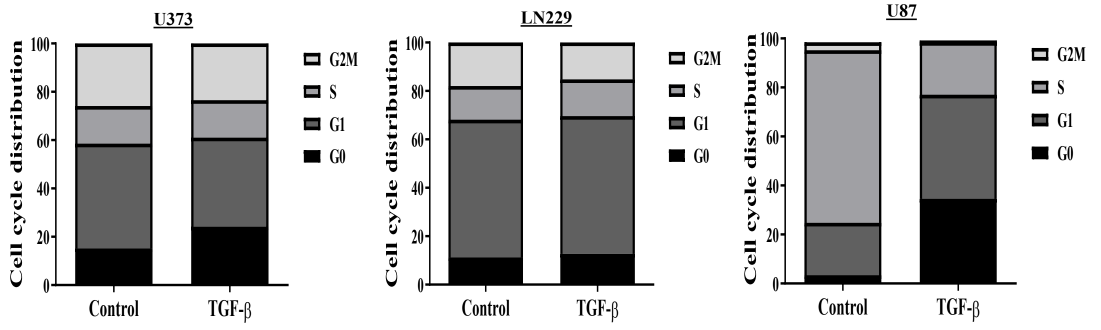


**(E)**

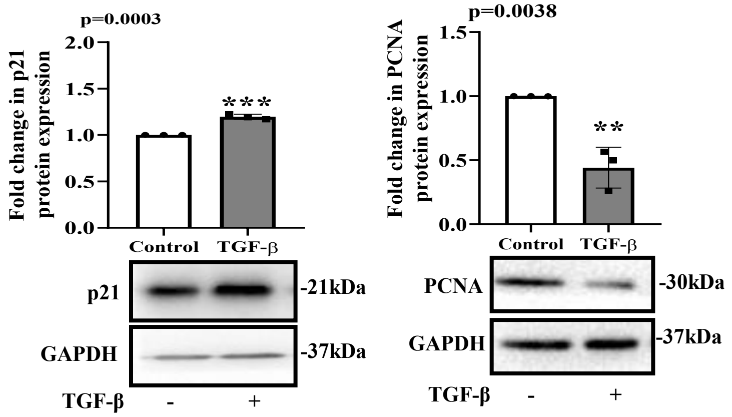


**(F)**

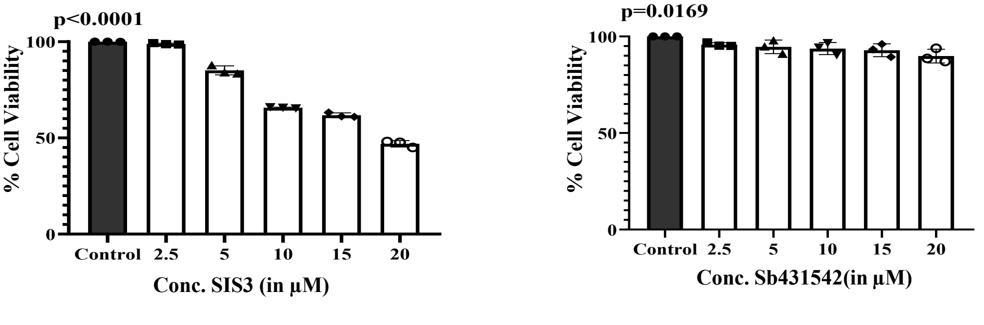

### Supplementary Figure 2

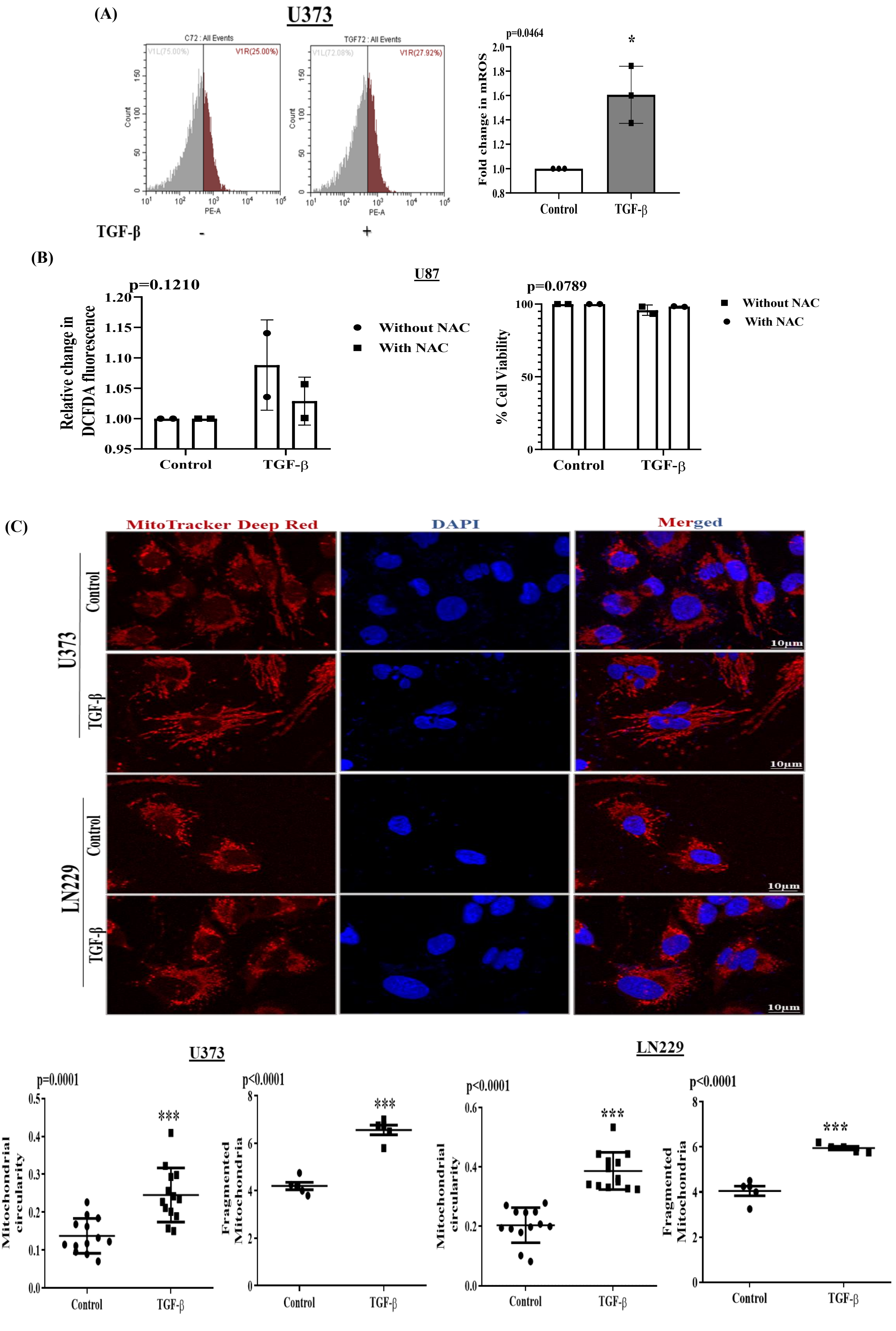

**Supplementary Figure 3**

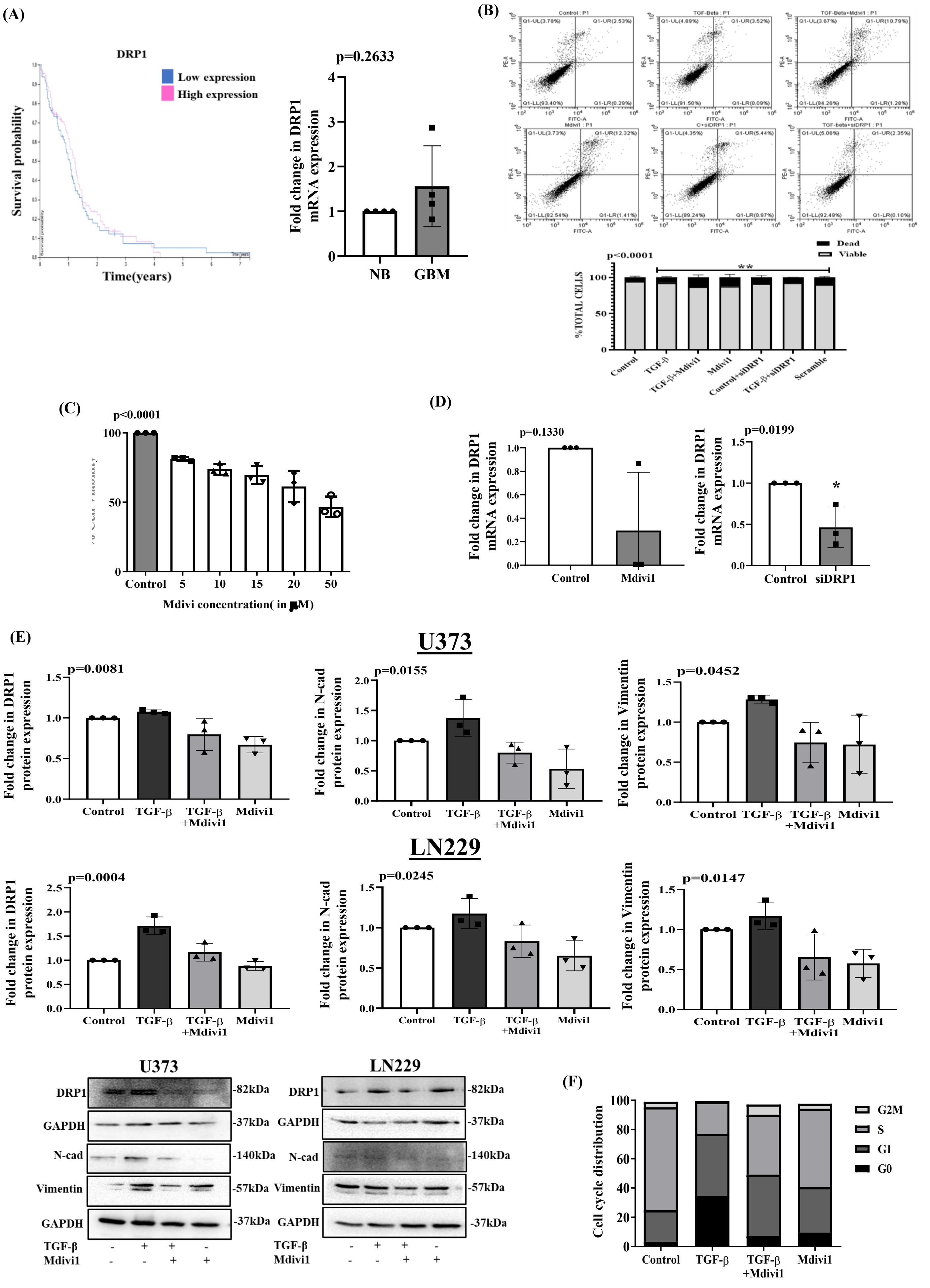
